# Glia of *C. elegans* coordinate the heat shock response independent of the neuronal thermosensory circuit and serotonin

**DOI:** 10.1101/2022.01.31.478522

**Authors:** Holly K. Gildea, Phillip A. Frankino, Sarah U. Tronnes, Corinne L. Pender, Hyun Ok Choi, Tayla D. Hunter, Shannon S. Cheung, Ashley E Frakes, Edward Sukarto, Andrew Dillin

## Abstract

As organisms age, they lose the ability to induce appropriate stress responses, becoming vulnerable to protein toxicity and tissue damage. Neurons can signal to peripheral tissues to induce protective organelle-specific stress responses. Recent work has demonstrated a novel and independent role of glia in inducing such responses. Here, we show that overexpression of heat shock factor 1 (*hsf-1*) in the four astrocyte-like cephalic sheath cells of *C. elegans* is sufficient to induce a non-cell autonomous cytosolic unfolded protein response (UPR), also known as the heat shock response (HSR), in distal cells. These animals upregulate the HSR in peripheral cells and have increased lifespan and resistance to heat stress. This glial HSR regulation is independent of the canonical neuronal thermosensory circuit and of known neurotransmitters but is dependent on the small clear vesicle release protein UNC-13. Additionally, HSF-1 and the FOXO transcription factor DAF-16 are partially required in peripheral tissues for increase of non-autonomous HSR, lifespan, and thermotolerance. We find that cephalic sheath glial *hsf-1* over-expression leads to increased pathogen resistance, which suggests a role for this signaling pathway in immune function.

## Introduction

Cellular insults occur as animals age that can cause dysfunction. Cells have compartment-specific signaling pathways that detect such insults, temporarily limit protein production, and upregulate protective genes, such as the protein folding assistant chaperones, to rescue cells from potentially toxic protein misfolding. As organisms experience damage over a lifetime, the cellular ability to mount responses to stress also declines (*1*, *2*). The process of aging perturbs cellular homeostasis by reducing organelle-specific unfolded protein response (UPR) induction and efficacy (*1*, *2*). Rescue of UPR functions by overexpression of activators in the nervous system increases healthspan and lifespan, indicating that UPRs are a potential therapeutic target for aging (*2*–*4*).

The compartment-specific UPR initiated by proteotoxic stress in the cytosol is known as the heat shock response (HSR) and is primarily mediated by the highly conserved transcription factor heat shock factor 1 (HSF-1) (*5*). Under non-stressed conditions chaperones such as HSP-70 and HSP-90 bind HSF-1, suppressing its activation (*6*). Upon detection of misfolded proteins in the cytosol the chaperones are titrated away from HSF-1, freeing the transcription factor to trimerize and translocate into the nucleus (*6*). There, HSF-1 upregulates chaperones and other genes that help resolve stress. HSF-1 activity declines with age, and this dysfunction occurs concomitant with worsening of cytosolic protein aggregation (*1*, *7*, *8*).

Recent work has established a unique role for the nervous system in initiating UPRs, including the HSR, across the whole organism (*2*–*4*, *9*, *10*). When the 302 neurons of *C. elegans* over-express *hsf-1*, animals exhibit a non-cell autonomous activation of the HSR in peripheral tissues, which leads to an increase in thermotolerance and lifespan (*3*). In *C. elegans*, heat sensing occurs via the canonical thermosensory circuit including AFD, AIY, and serotonergic neurons, and is required for behaviors such as thermotaxis (*11*). Interestingly, electrical activation of these cells, including AFD sensory neurons and downstream ADF serotonergic neurons, has been shown to induce peripheral HSF-1 activation in addition to canonical heat sensing behaviors (*12*, *13*). Thus, neural activity due to the sensory experience of heat, a potentially damaging insult, is coupled to the relevant organismal intracellular heat shock stress response.

Despite neuronal ability to induce the HSR non-cell autonomously when exogenously activated, neurons are likely not the most potent responders of the nervous system (*7*, *8*, *14*, *15*). Hyperthermia induces chaperone expression in neural cells; however, glia, particularly astrocytes, upregulate chaperones more robustly than neurons in these conditions (*14*, *15*). *In vitro* data suggest that glia may even provide chaperones to neurons directly (*16*). Neurons also aberrantly degrade HSF1 in several neurodegenerative disease conditions, including Alzheimer’s and Huntington’s Diseases, in model organisms and human tissue (*7*, *8*). These findings suggest that glia, not neurons, are likely the primary coordinators of cytosolic stress responses in the nervous system.

*C. elegans* glia play an important role in regulation of cellular stress and longevity (*17*, *18*). The 56 glia of *C. elegans* perform classic glial functions, supporting neuronal development, participating in synapses, and providing neurotransmitter and metabolic support to neurons (*19*, *20*). Four of these cells, the cephalic sheath (CEPsh) glia, most closely resemble mammalian astrocytes (*19*). CEPsh glia are poised at a unique junction of peripheral tissues and the nervous system. They ensheath processes of sensory neurons that project their endings into the environment. These glia also surround the nerve ring, forming a barrier between the nerve ring and the rest of the body (*20*). Recent work has demonstrated that these cells are able to induce organelle-specific stress responses non-cell autonomously in the case of the endoplasmic reticulum (ER) and mitochondria, but their role in cytosolic protein stress sensing and signaling has not been explored (*17*, *18*).

Here, we find that over-expression of *hsf-1* in the four CEPsh glia of *C. elegans* is able to coordinate an organismal HSR, confer stress resistance, and extend lifespan. Signaling of the glial HSR relies on a mechanism distinct both from that of neuronal HSR induction and from other glial stress responses. This response is independent of the canonical *C. elegans* neuronal thermosensory circuit and of dense core vesicle release. It requires the presence of small clear vesicle release machinery, though no single neurotransmitter known to be released through these vesicles is independently required for the peripheral HSR induction. CEPsh glial HSF-1 coordinates the upregulation of immune regulators, resulting in pathogen resistance. These data implicate *C. elegans* CEPsh glia as primary sensors and signalers of protein health insults, which can flexibly and specifically adopt signaling strategies to coordinate health and longevity across the organism.

## Results

To assess whether the four *C. elegans* CEPsh glia upregulate a protective HSR organismally in response to *hsf-1*, we created strains over-expressing *hsf-1* under the CEPsh glia-specific promoter *hlh-17 (hlh-17p::hsf-1)* (*17*, *21*, *22*). To evaluate the impact of *hlh-17p::hsf-1*, CEPsh glial *hsf-1*, on longevity, we first assayed lifespan under normal culture conditions. We found that CEPsh glial *hsf-1* animals were longer lived than wild type N2 worms (Figure 1A, Supp Fig 1A). We also observed that this coincides with a suppression of fecundity (Supp Fig 1B,C). This is consistent with existing work suggesting increased HSF-1 function in lifespan is a trade-off with reproductive fitness (*1*, *23*) This consequence may explain why HSF-1 expression is tightly titrated across evolution. We next examined heat stress tolerance and found that CEPsh glial *hsf-1* animals are robustly thermotolerant compared to wildtype N2 animals (Figure 1B, Supp Fig 1D). To test whether the increased longevity and stress tolerance of CEPsh glial *hsf-1* animals correlate with organismal induction of HSR genes, we used a fluorescent transcriptional reporter for *hsp-16.2*, a heat shock chaperone induced upon heat stress. Using this system, we found that CEPsh glial *hsf-1* animals strongly upregulated HSR chaperones upon heat stress compared to reporter worms alone, and that this increased expression was evident throughout the worm, predominantly visible in the intestine (Figure 1C and D, Supp Fig 1E,F). We thus determined that CEPsh glia non-autonomously induce the HSR, increasing longevity, stress response activation, and stress tolerance.

**Figure 1:**
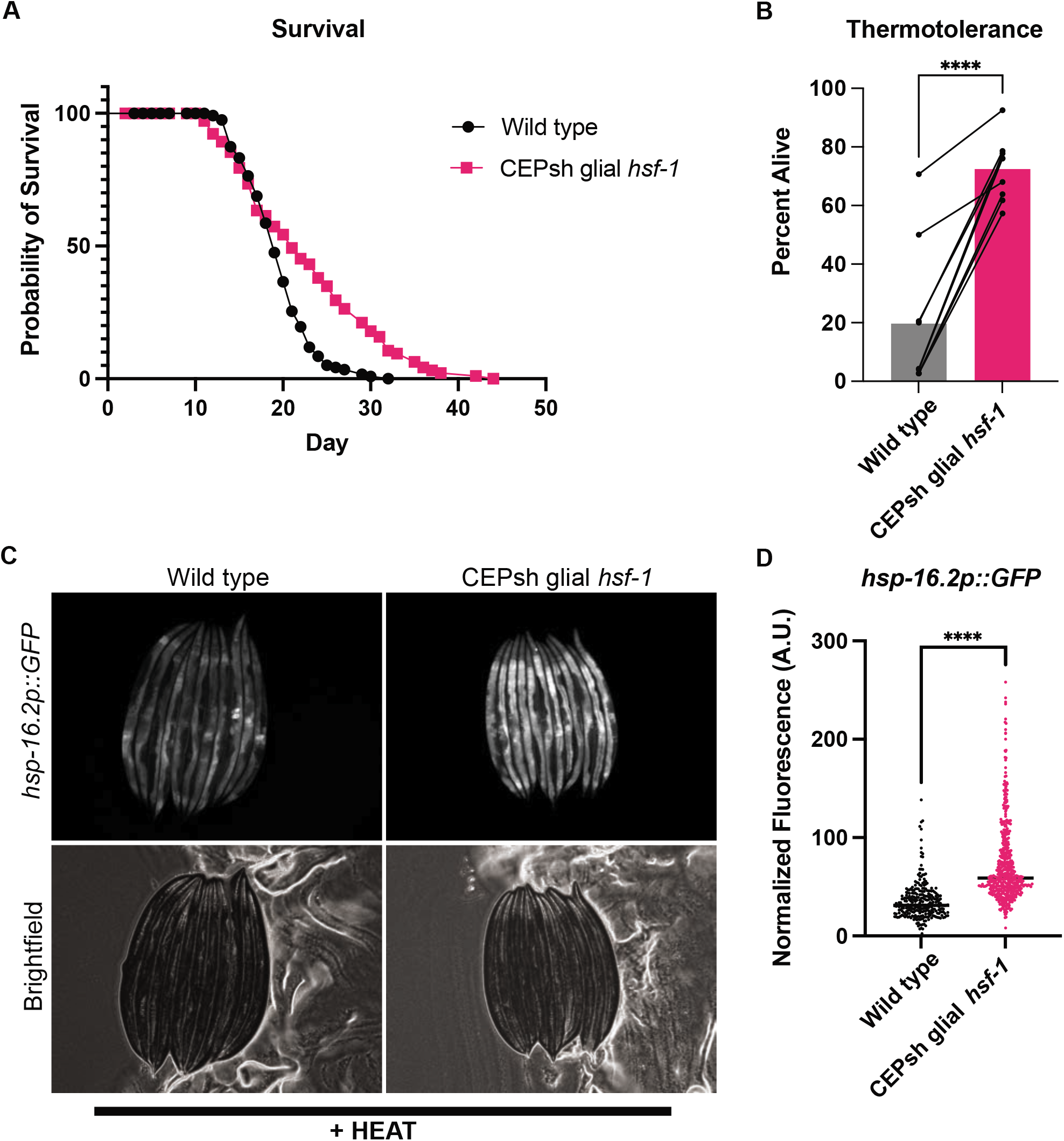
Overexpression of *hsf-1* in CEPsh glia increases lifespan and stress tolerance by non-autonomously inducing the heat shock response. A) Survival of *hlh-17p::hsf-1* (integrated array) is significantly greater than wild type N2 at 20C. Mean survival of N2 = 19 days, *Is1(hlh-17p::hsf-1)* = 21 days, p= 0.017. B) Thermotolerance of *Ex(hlh-17p::hsf-1)* array expressing animals is significantly increased relative to wild type N2 animals after heat stress at 34C for 12-16h. Connected points are individual experiments, p<0.0001. C) *hsp-16.2p::GFP* transcriptional reporter worms with and without *Ex(hlh-17p::hsf-1)* after mild heat stress and recovery, lined up head to tail. D) Quantification of worm fluorescence strains in C in large particle flow cytometry via COPAS biosorter. *Ex(hlh-17p::hsf-1)* worms are significantly brighter, p < 0.0001.

Under natural heat sensing conditions, the AFD thermosensory neuron signals to the AIY interneuron, which is upstream of serotonergic neurons such as NSM and ADF(Figure 2A) (*12*, *24*). To ascertain whether the heat stress response to glial *hsf-1* is mediated by the canonical thermosensory neuronal circuitry, we measured induction of *hsp-16.*2 in mutants defective in AIY interneuron formation, *ttx-3(ks5)*, with CEPsh glial *hsf-1* over-expression. *ttx-3(ks5)* mutant animals have been previously shown to decrease induction of the HSR in otherwise wild type animals (*9*). We found that under acute heat shock, *ttx-3(ks5)* mutant animals failed to suppress the increased levels of *hsp-16.2* due to CEPsh glial *hsf-1* (Figure 2B,C). These data suggest that the contribution of glia to HSR induction must be either downstream of or independent of AIY thermosensory neuron function, unlike the neuronal HSR.

**Figure 2:**
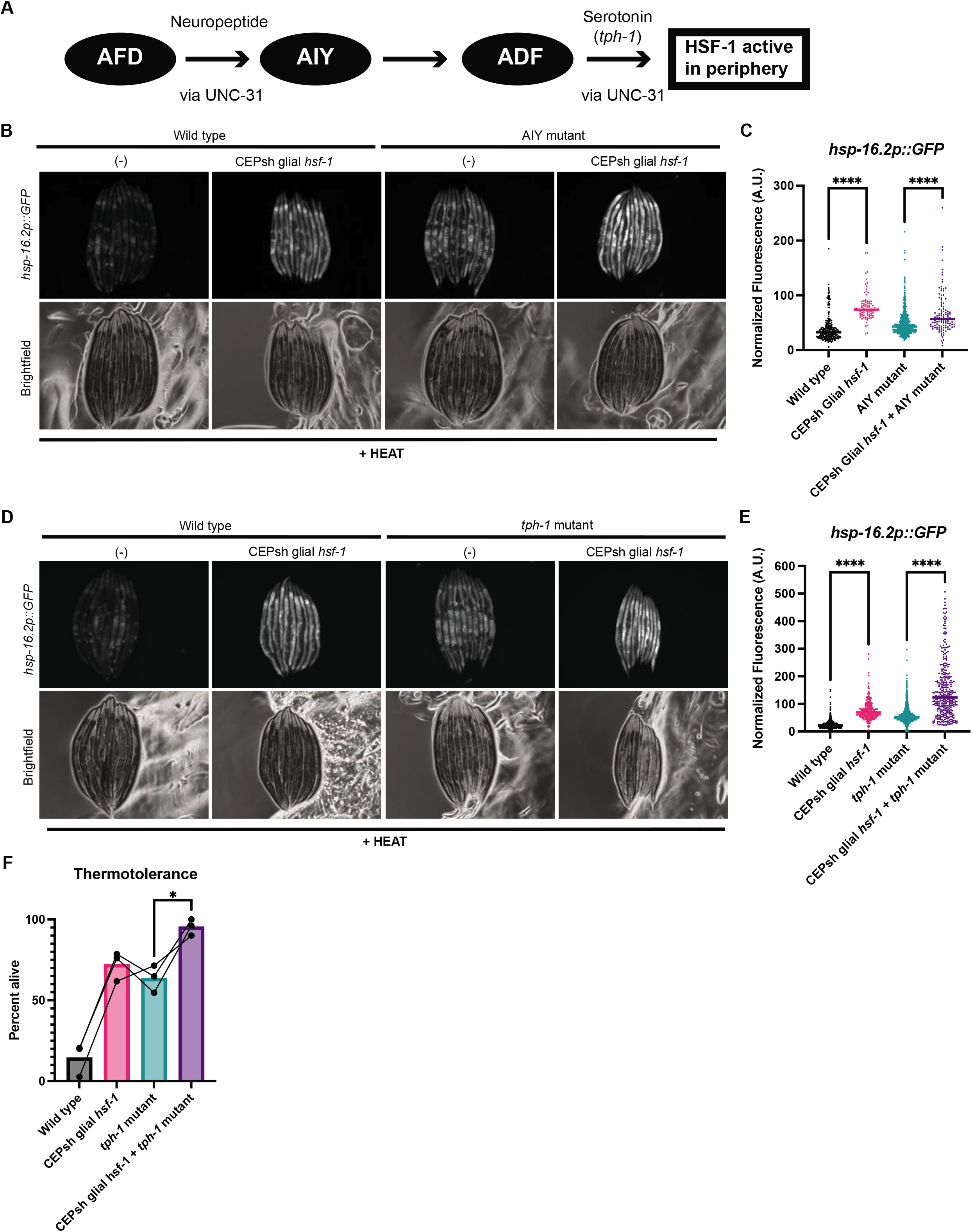
The canonical thermosensory circuit is dispensable for CEPsh glial *hsf-1* signaling. A) Schematic of the thermosensory circuit and relevant signaling components. B) Mutants for AIY, *ttx-3(ks5)*, with and without *Ex(hlh-17p::hsf-1)* imaged for the *hsp-16.2p::GFP* after mild heat stress and recovery. *Ex(hlh-17p::hsf-1); ttx-3(ks5)* exhibit higher *hsp-16.2p::GFP* fluorescence than *ttx-3(ks5)* alone. C) Quantification of strains in B via large particle flow cytometry. *Ex(hlh-17p::hsf-1); ttx-3(ks5)* are brighter than *ttx-3(ks5)* alone (p < 0.0001). D) Mutants for tryptophan hydroxylase, *tph-1(mg280)*, with and without *Ex(hlh-17p::hsf-1)* imaged for the *hsp-16.2p::GFP* after mild heat stress and recovery. *Ex(hlh-17p::hsf-1); tph-1(mg280)* animals exhibit higher *hsp-16.2p::GFP* fluorescence than *tph-1(mg280)* animals alone. E) Quantification of strains in D via large particle flow cytometry. *Ex(hlh-17p::hsf-1); tph-1(mg280)* animals are significantly brighter than *tph-1(mg280)* alone (p < 0.0001). F) Thermotolerance of wild type N2 animals, *Ex(hlh-17p::hsf-1)* animals, *tph-1(mg280)* animals, and *Ex(hlh-17p::hsf-1); tph-1(mg280)*. Data points are individual experiments, and connecting line indicates paired trials. Error bars are S.D. Thermotolerance of *Ex(hlh-17p::hsf-1); tph-1(mg280)* is significantly greater than *tph-1(mg280)* (p=0.04).

Due to the apparent divergence of glial HSR regulation from this canonical thermosensory circuit component, we next asked whether other members of this core circuitry were dispensable for CEPsh glial *hsf-1* signaling. Serotonin and serotonin receptor activity are also required for downstream sensing of the AFD/AIY thermosensory circuit (*12*). Therefore, we next examined animals with the *tph-1(mg280)* mutation, which lack functional tryptophan hydroxylase and are unable to synthesize serotonin, for induction of *hsp-16.2* by glial *hsf-1*. We found that serotonin synthesis is dispensable for peripheral induction of *hsp-16.2* by CEPsh glial *hsf-1* activation, implying that glial HSR induction is independent of serotonin (Figure 2D,E). To further examine the role of serotonin, we subjected CEPsh glial *hsf-1* animals with and without the *tph-1(mg280)* mutation to chronic heat stress. We found that serotonin synthesis is not required for the glial *hsf-1*-mediated increase in thermotolerance (Figure 2F). Taken together, the signaling of the HSR by CEPsh glia does not occur via the canonical neuronal heat stress pathway, nor does it require serotonin.

We next asked which signaling molecules might be responsible for this non-cell autonomous signaling of the HSR to peripheral tissues by CEPsh glia, if not serotonin. Previous studies of stress signaling from glia in the case of the ER and mitochondrial UPRs implicated neuropeptides (*17*, *18*). CEPsh glia could regulate distinct stress responses with similar or distinct signals. Dense core vesicles are required for the release of larger cargos, like neuropeptides, while small clear vesicles are required for neurotransmitter release (*25*, *26*). To determine whether small clear vesicles or dense core vesicles might be required for non-cell autonomous HSR induction in CEPsh glial *hsf-1* animals, we used mutants for the vesicular release components *unc-13* and *unc-31*, respectively (*25*, *26*). We found that loss of small clear vesicle fusion via *unc-13(s69)* suppressed CEPsh glial *hsf-1* non-cell autonomous induction of *hsp-16.2* (Figure 3A, B). In contrast, loss of dense core vesicle fusion via *unc-31(e958)* mutation failed to suppress the increase in peripheral *hsp-16.2* activation (Figure 3C,D) (*25*, *26*). Taken together, these data indicate that CEPsh glial signaling of the HSR relies on cargo enclosed in small clear vesicles and is independent of dense core vesicle neuropeptide signaling, unlike the neuronal and glial mitochondrial UPR and glial UPR ER responses (*17*, *18*).

**Figure 3:**
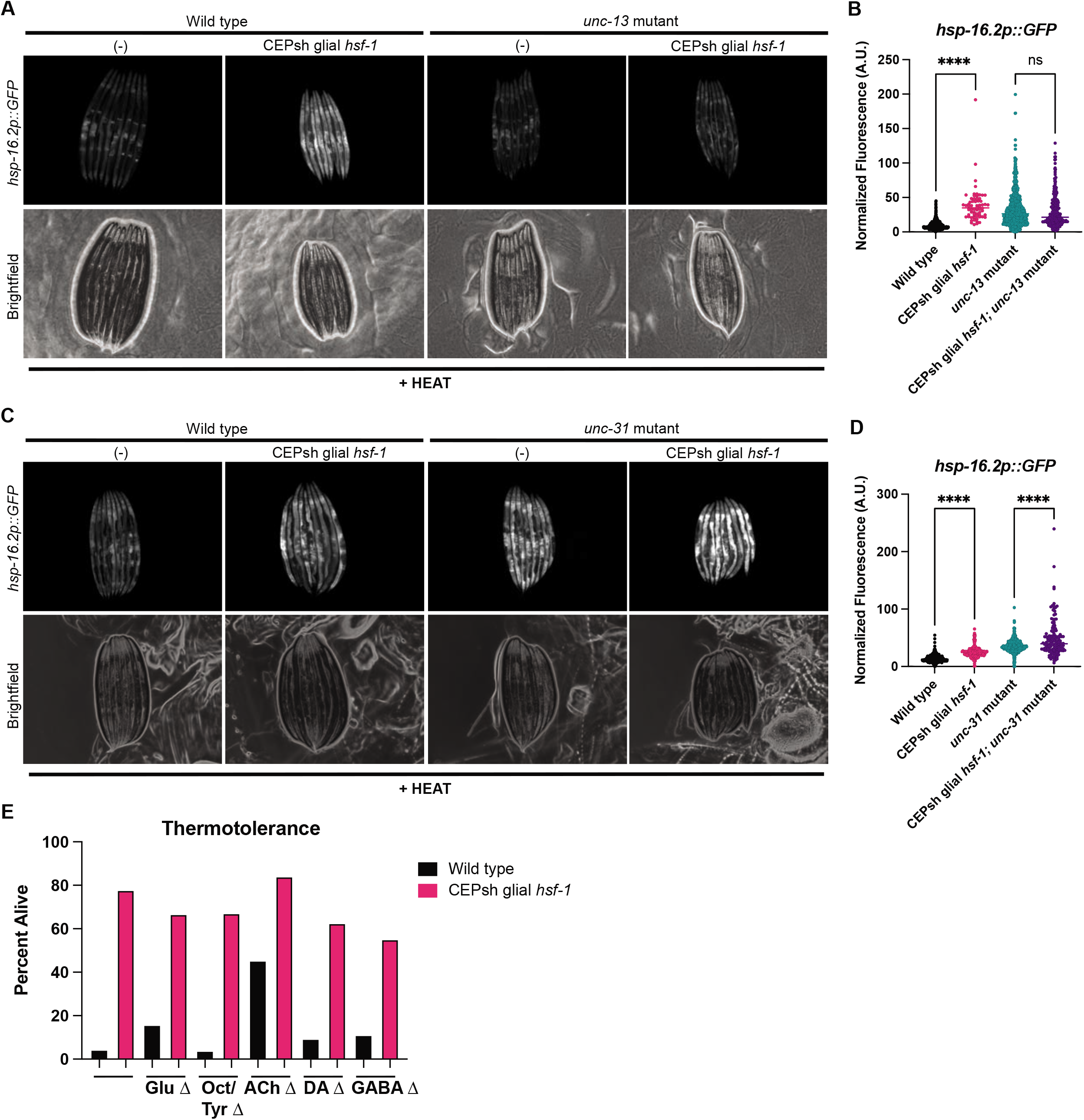
Small clear vesicles, but not dense core vesicles, are required for CEPsh glial *hsf-1* signaling via an unknown cargo. A) Mutants for small clear vesicle release, *unc-13(s69)*, with and without *Ex(hlh-17p::hsf-1)* imaged for *hsp-16.2p::GFP* after mild heat stress and recovery. *Unc-13(s69)*, *Ex(hlh-17p::hsf-1)* animals have no increase in *hsp-16.2p::GFP*. B) Quantification of strains in A via large particle flow cytometry. Fluorescence of *Ex(hlh-17p::hsf-1); unc-13(s69); hsp-16.2p::GFP* animals is not significantly increased vs. *unc-13(s69); hsp-16.2p::GFP*. C) Mutants for dense core vesicle release, *unc-31(e958)*, with and without *Ex(hlh-17p::hsf-1)* imaged for the reporter *hsp-16.2p::GFP* after mild heat stress and recovery. *Unc-31(e958)*, *Ex(hlh-17p::hsf-1)* animals exhibit an increase in *hsp-16.2p::GFP*. D) Quantification of strains in C via large particle flow cytometry. Fluorescence of *Ex(hlh-17p::hsf-1); unc-31(se958); hsp-16.2p::GFP* animals is significantly increased relative to *unc-31(e958); hsp-16.2p::GFP* (p<0.0001). D) Thermotolerance of *Ex(hlh-17p::hsf-1)* (pink) and wild type N2 (black) animals. Neurotransmitter mutation is on the x axis, such that the left-most comparison of *Ex(hlh-17p::hsf-1)* and wild type N2 is in the wild type background, followed by Glu = *eat-4(ky5)*, Oct/Tyr = *tdc-1(n3419)*, ACh = *unc-17(e245)*, DA = *cat-2(n4547)*, GABA = *unc-25(e156)*. Representative experiment displayed, with at least 3 replicates for each mutant condition, less *unc-25* with 2 replicates.

As the canonical cargoes for small clear vesicles in the worm are neurotransmitters, we selected a set of mutants in synthesis or vesicular loading for each of the known neurotransmitters in *C. elegans*. Having already assessed serotonin, we turned our attention to mutants defective in signaling by glutamate (*eat-4*), GABA (*unc-25*), dopamine (*cat-2*), acetylcholine (*unc-17*), and octopamine and tyramine (*tdc-1*). Using the thermotolerance assay, we found that the increase in survival due to glial *hsf-1* is preserved in the absence of dopamine, GABA, and octopamine/tyramine (Figure 3E, Supp. Figure 2). Further, there is a trend towards significance in the survival increase of acetylcholine and glutamate mutants (Figure 3E, Supp. Figure 2). Notably, acetylcholine and glutamate are canonical small clear vesicle cargoes, though the apparent partial suppression of effect suggests neither transmitter is independently required for signaling (Supp. Figure 2D,E). These data suggest that no known neurotransmitter is independently responsible for organismal protection against heat stress conferred by glial *hsf-1*, though the signal is likely contained in an UNC-13-mediated vesicle.

Neuronal HSR signaling relies on both HSF-1 and the insulin signaling-related FOXO homolog DAF-16 in peripheral tissues to enact survival benefits. Therefore, we tested the requirement for these transcription factors in peripheral HSR activation of CEPsh glial *hsf-1* animals (*3*). Neurons and glia of *C. elegans* are partially resistant to RNA interference (RNAi), allowing us to directly interrogate peripheral signaling requirements. Examining induction of the *hsp-16.2* transcriptional reporter, we found that, as predicted, *hsf-1* is required in peripheral cells for most HSR chaperone induction (Figure 4A, B). In contrast, *daf-16* is not required for *hsp-16.2* induction (Figure 4C, D). Despite these differences in chaperone induction, the increase in lifespan due to CEPsh glial *hsf-1* was largely dependent on both *hsf-1* and *daf-16* in peripheral tissues (Figure 4E-G). Further, *hsf-1* and *daf-16* seem to be at least partially required for thermotolerance increase (Supp. Figure 3A,B). Overall, these data indicate that *hsf-1* and *daf-16* act in concert to regulate the protective phenotypes of CEPsh glial *hsf-1* animals in the peripheral tissues.

**Figure 4:**
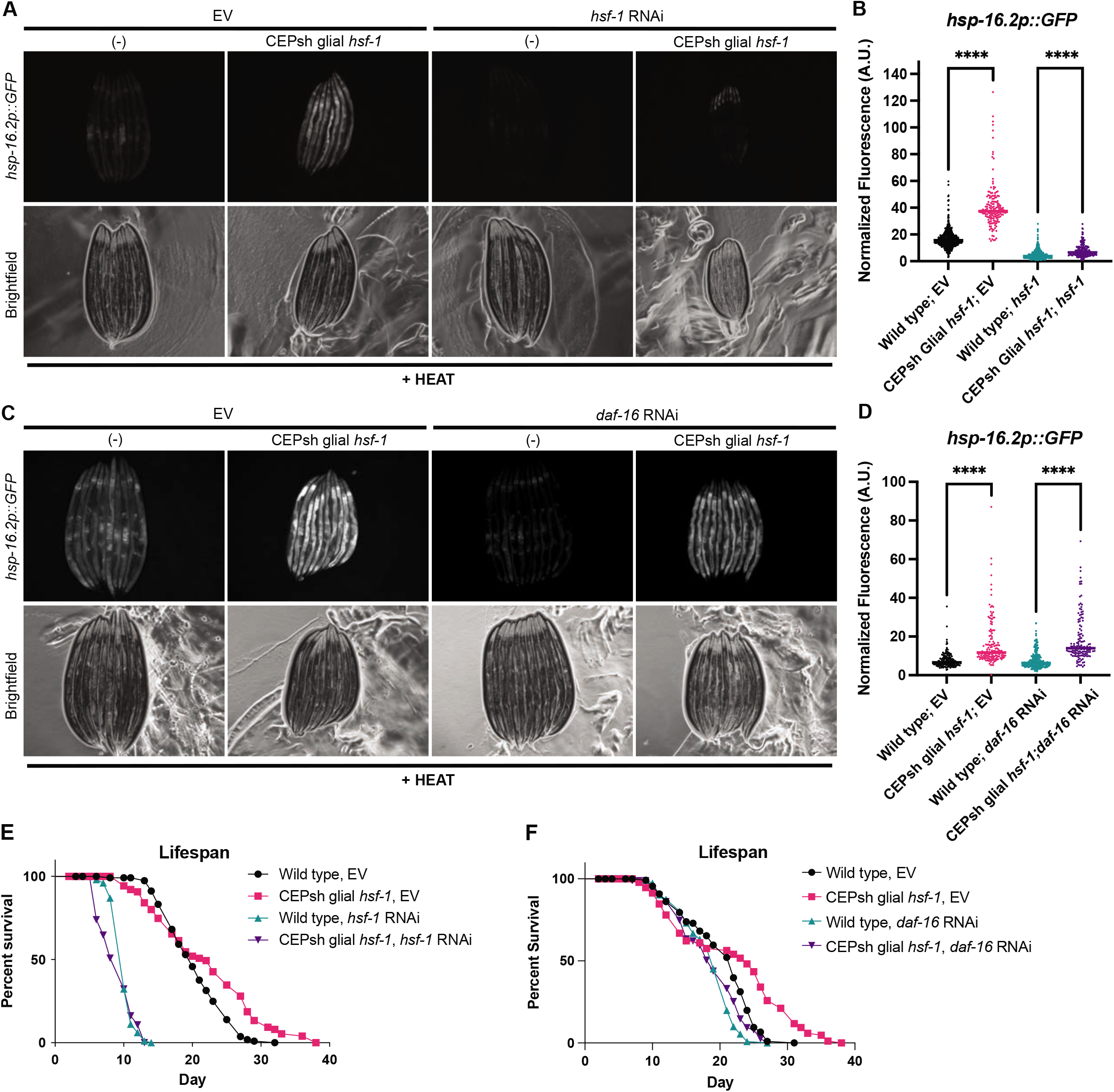
HSF-1 and DAF-16 are required for benefits of CEPsh glial *hsf-1* in peripheral tissues. A) Wild type and *Ex(hlh-17p::hsf-1)* imaged for the reporter *hsp-16.2p::GFP* after mild heat stress and recovery under empty vector (EV) and *hsf-1* RNAi conditions. RNAi against *hsf-1* suppresses *hsp-16.2p::GFP*. B) Quantification of strains in A via large particle flow cytometry. *Ex(hlh-17p::hsf-1)*-related *hsp-16.2p::GFP* induction is partially suppressed by *hsf-1* RNAi, though *Ex(hlh-17p::hsf-1)* on *hsf-1* RNAi remains significantly increased vs. wild type N2 on *hsf-1* RNAi (p<0.0001). C) Wild type and *Ex(hlh-17p::hsf-1)* imaged for *hsp-16.2p::GFP* after mild heat stress and recovery on EV and *daf-16* RNAi. D) Quantification of strains in C via large particle flow cytometry. *Ex(hlh-17p::hsf-1)* on *daf-16* RNAi remains significantly increased vs. wild type N2 animals on *daf-16* RNAi (p<0.0001). E) Lifespan of *Is1(hlh-17p::hsf-1)* vs. wild type N2 worms on EV and *hsf-1* bacteria. Mean survival of N2 on EV = 20 days, *Is1(hlh-17p::hsf-1)* on EV = 22 days, N2 on *hsf-1* RNAi = 10 days, and *Is1(hlh-17p::hsf-1)* on *hsf-1* RNAi = 10 days, p<0.0001. F) Lifespan of *Is1(hlh-17p::hsf-1)* vs. wild type N2 worms on EV and *daf-16* bacteria. Mean survival of N2 on EV = 22 days, *Is1(hlh-17p::hsf-1)* on EV = 24 days, N2 on *daf-16* RNAi = 19 days, and *Is1(hlh-17p::hsf-1)* on *daf-16* RNAi = 19 days, p= 0.0012.

Beyond known HSR effectors, we next sought to identify organismal changes in gene expression that might shed light on peripheral tissues’ interpretation of glial *hsf-1* signaling. Whole animal RNA sequencing (RNA-seq) revealed substantial gene expression changes in CEPsh glial *hsf-1* animals compared to wild type N2 animals, with 692 genes significantly upregulated and 272 genes downregulated (adj-p <= .05 and log2(FC) of greater than 1 or less than −1, respectively). In CEPsh glial *hsf-1* animals, *hsf-1* is significantly upregulated, and HSR proteins *hsp-16.2* and *hsp-70* are mildly increased, while chaperones for the ER and mitochondrial UPRs were either unchanged or downregulated, respectively (Figure 5A). To identify high confidence *hsf-1*-regulated genes that were differentially expressed in the CEPsh glial *hsf-1* animals, we generated a list of genes that were previously reported to be *hsf-1* targets and had HSF-1 binding sites in the immediate upstream region from the start codon (*27*). Indeed, many high confidence HSF-1 target genes were significantly upregulated or downregulated (p<0.05) in CEPsh glial *hsf-1* animals (Figure 5B).

**Figure 5:**
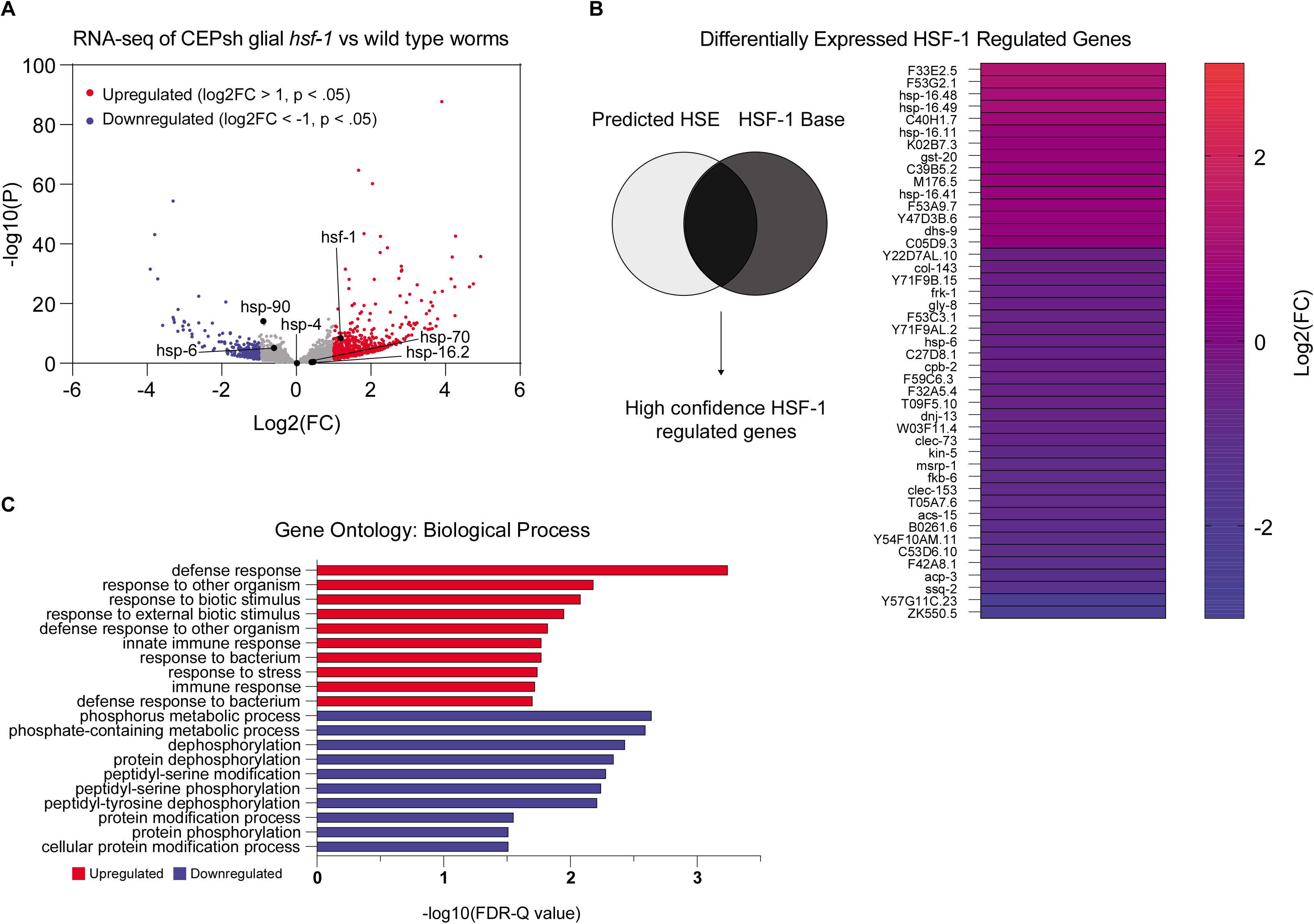
CEPsh glial *hsf-1* induces peripheral changes in HSF-1-regulated genes and the immune response. A) Volcano plot demonstrating magnitude (Log2(FC)) and significance (−log10(p-value)) of changes in gene expression from whole-animal RNA sequencing of *Is1(hlh-17p::hsf-1)* versus wild type N2. Labeled genes are stress genes, including *hsf-1* and HSR chaperones *hsp-70*, *hsp-16.2* (HSF-1 regulated) and *hsp-90* (non-HSF-1 regulated), as well as ER UPR chaperone *hsp-4* and mitochondrial UPR chaperone *hsp-6*. B) High confidence HSF-1-regulated genes are displayed alongside their Log2(FC) values, color coded from cool (down-regulated) to warm (up-regulated). C) The top ten Gene Ontology (GO) terms for up- (red) and down- (blue) regulated genes are displayed with their −log10 corrected FDR-Q value.

To evaluate the categories in which whole animal gene expression was altered by sensing CEPsh glial *hsf-1*, we used gene ontology (GO) analysis of upregulated and downregulated genes (*28*, *29*). GO term enrichment analysis of the significantly upregulated genes contained GO terms concerning the immune response and stress responses generally, while GO terms associated with the significantly downregulated genes highlighted protein modification (Figure 5C). Overall, sequencing analysis of the CEPsh glial *hsf-1* animals reveals a broad upregulation of immune and stress response genes with differential expression of many *bona fide* HSF-1 target genes.

Infectious insults are important natural environmental stimuli for worms, and infection is a major cause of death across the organism’s lifespan (*30*). The bacteria *Pseudomonas aeruginosa* is pathogenic to *C. elegans*, and *hsf-1* is required for normal survival on *P. aeruginosa* (*31*). Further, heat shock chaperones are activated upon exposure to the bacteria (*13*). Given data suggesting a broad upregulation of immune genes in CEPsh glial *hsf-1* animals, we hypothesized that CEPsh glial *hsf-1* might induce pathogen resistance. We therefore tested resistance of CEPsh glial *hsf-1* versus wild type N2 worms on the *P. aeruginosa* PA14 strain using the slow killing assay and found a robust increase in survival in CEPsh glial *hsf-1* animals (Figure 6A). These data suggest that CEPsh glial upregulation of HSF-1 activity drives a *bona fide* immune response that protects the animals from bacterial infection.

**Figure 6:**
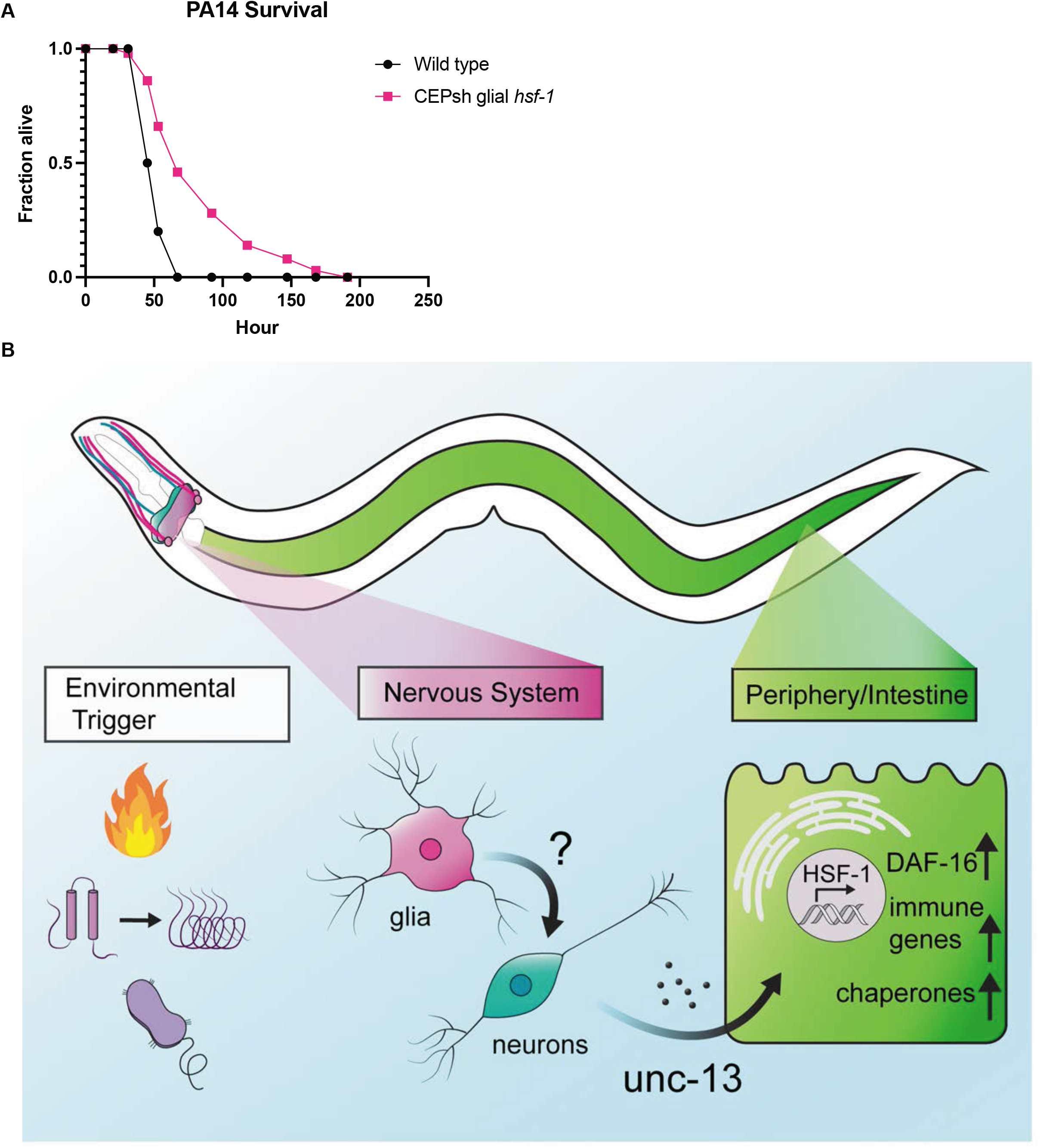
CEPsh glial *hsf-1* induces immune resistance. A) Survival of *Ex(hlh-17p::hsf-1)* is increased versus wild type N2 animals on *Pseudomonas aeruginosa* bacteria. Mean survival for wild type N2 animals = 53h, for *Ex(hlh-17p::hsf-1)* animals = 67h. p < 0.0001 B) Schematic summarizing findings. Heat, immune challenge, or misfolded proteins may activate signaling. Glial activation of *hsf-1* likely signals to neurons, causing release of some *unc-13*-dependent cue to peripheral cells. Downstream cells activate HSF-1 and DAF-16 as well as immune factors, to increase lifespan, stress tolerance, and immune resistance.

## Discussion

We have identified a unique role for the four astrocyte-like CEPsh glia of *C. elegans* in coordinating a non-cell autonomous heat shock response. Animals over-expressing *hsf-1* in CEPsh glia are both more tolerant to heat stress and longer-lived. These phenotypes correlate with an increase in HSR chaperones across the animal, demonstrating the ability of CEPsh cells to induce the stress response in distal tissues by a diffuse signaling mechanism.

*C. elegans* have 959 somatic cells, 302 neurons, and 56 glia, among which only four are CEPsh glia. Previous work demonstrated beneficial effects on longevity when over-expressing *hsf-1* in all 302 neurons, which amount to nearly one third of the animal’s cells; in this study, however, we over-express *hsf-1* in fewer than 0.5% of all cells of the worm and find similar effects on lifespan and stress tolerance (*3*). These data suggest that the worm is particularly responsive to signaling from CEPsh glia. As glia are best known for their interactions with neurons, it also suggests that glial stress responses play a larger role in regulating neuronal stress response activity.

Interestingly, glial coordination of the HSR is independent of known components of the canonical thermosensory circuit. The AIY interneuron has been previously identified as a hub for integrating heat sensing (*3*, *9*); however, we find that this neuron is dispensable for the CEPsh glial HSR. Our identification of an UNC-13-dependent, non-AIY mediated glial signal for regulating lifespan via *hsf-1* signaling implies that two distinct pathways may be at work: one composed of thermosensory neuron components, and the other downstream of CEPsh glia. This is further evident in the independence of glial HSR signaling from serotonin, a downstream component of the thermosensory circuit.

Notably, the serotonin-independent nature of the CEPsh glial HSR mechanism differentiates it not only from the neuronal circuit controlling the HSR, but also from other neuronally-controlled proteostasis responses. Neurons ostensibly converge to regulate protein homeostasis using serotonin, which has been previously implicated in neuronal regulation of stress resistance via the HSR (*12*), the mitochondrial UPR (*32*), and the endoplasmic reticulum UPR (*33*). The CEPsh glial HSR mechanism is thus novel not only in the context of HSR regulation, but also in neural regulation of UPRs broadly. Furthermore, lifespan regulation by CEPsh glial signaling in the cases of the ER (*34*) and mitochondria (*18*) rely on neuropeptides, which are released via dense core vesicles, whereas the CEPsh glial HSR functions independent of dense core vesicle release. Although neuropeptides are certainly powerful diffusible cues, the HSR data suggest release of neuropeptides is not the automatic glial response to stress across circumstances. Rather, the cells are able to flexibly react to internal states to induce specific programs in distal cells.

We find a requirement for the small clear vesicle release protein UNC-13 in the non-cell autonomous communication of the HSR by CEPsh glia; however, the identity of the cue or cues contained in such vesicles remains unclear. As UNC-13 is thought to be expressed specifically in neurons, the most likely model involves CEPsh glial recruitment of neurons for signaling. We genetically disrupted production or packaging of serotonin, dopamine, octopamine/tyramine, acetylcholine, GABA, and glutamate, and failed to see a robust reduction in HSR signaling. Therefore, these signals could be functioning redundantly to induce the response, another unidentified cargo may be loaded into small clear vesicles, or glia may be modulating neurons in some other way, for example, by reuptake of neurotransmitters. Firstly, a combination of transmitters may signal the glial HSR, potentially both acetylcholine and glutamate, for example. Also, distinct non-canonical cues may be at play. A novel small clear vesicle cargo derived from glia or from neurons may be responsible for this signaling. Several distinct stressors have been shown to induce lipids as a glial-neuronal signal, for example, which this work cannot rule out (*35*–*37*). CEPsh glia may also alter neurotransmitter release via damage signals, immune molecules, or even chaperones themselves, though these mechanisms are not well described in the glia of *C. elegans*. Finally, CEPsh glia previously have been shown to alter neuronal activity via neurotransmitter reuptake, particularly in the case of glutamate (*38*). CEPsh glial *hsf-1* may induce such modulation of neuronal activity and could feasibly accomplish non-cell autonomous HSR signaling through these means.

Despite substantial differences in initiation, both the glial and neuronal HSRs converge on the peripheral factors HSF-1 and DAF-16. The whole animal RNA sequencing data presented in this study suggest that *hsf-1* may be transcriptionally upregulated in response to the glial HSR. Less surprisingly, peripheral HSF-1 seems to be required for induction of HSR chaperones, implying activation of the transcription factor’s canonical activity in non-glial cells is necessary for the protein homeostasis effects of CEPsh glial *hsf-1*. The beneficial effects of glial *hsf-1* on lifespan are wholly dependent on *hsf-1*, in contrast to the neuronal *hsf-1* model in which HSF-1 is only partially required (*3*). These data suggest that HSF-1 may be an upstream component of the peripheral response, which may be able to activate other beneficial factors downstream. By contrast, the FOXO transcription factor DAF-16, which has been previously implicated in lifespan extension across perturbations including the neuronal HSR, is only partially required for glial *hsf-1* phenotypes (*3*). DAF-16 is canonically repressed by kinases downstream of the insulin receptor DAF-2 as part of the insulin and IGF-1 signaling (IIS) pathway, and activation of DAF-16 is generally correlated with an increase in longevity (*39*–*41*). In the CEPsh glial *hsf-1* paradigm, DAF-16 is at least partially required for lifespan extension, thermotolerance and, to a lesser extent, chaperone induction. However, in all cases a slight increase remains despite peripheral knockdown of *daf-16*, supporting the hypothesis that DAF-16 may be downstream of HSF-1 or other induced protective factors.

We unbiasedly evaluated whole animal gene expression by RNA sequencing and found a surprising enrichment of immune-related genes upregulated in CEPsh glial *hsf-1* animals. HSF-1 has been previously implicated in immune function, and its role in pathogen resistance is independent of the canonical PMK-1/MAPK innate immune pathway, instead operating in a chaperone-dependent manner (*31*, *42*). In the worm, infection is a major cause of death, detectable by pharyngeal swelling, and *hsf-1* knockdown increases pharynx bacterial colonization (*30*, *43*). Data here indicate that CEPsh glia are able to induce a pro-immune and pro-longevity program by activating *hsf-1*, possibly increasing cellular protection from pathogens via induction of chaperones and immune response genes. Interestingly, by activating HSF-1-related genes specifically in this paradigm we were able to achieve effective increase of immune function in adult animals without a deleterious effect of prolonged immune activation on longevity. If increased HSF-1 function can both protect cells from proteotoxicity and pathogenic insults, we would anticipate that its activity would be positively selected evolutionarily. However, the negative impact of *hsf-1* upregulation on reproductive function demonstrated here suggests evolutionary titration of function balances these phenotypes to preserve the health of parents and offspring.

CEPsh glia are well positioned to receive cues from the environment, neurons, and peripheral tissues. This study, along with those detailing the role of these cells in the ER and mitochondrial UPRs, suggests that these cells may act as sensory organs particularly for organismal insults, inducing relevant and specific stress responses across the whole animal (*18*, *34*). The worm has no circulating adaptive immune system; however, the nervous system of *C. elegans* serves as an immune effector, regulating responses to toxic stimuli in both behavior and cellular programs. The connection between nervous system function and immune signaling in this case points to the larger role of the nervous system itself as the prototype for adaptive immunity.

CEPsh glia are thus able to coordinate protective functions by non-cell autonomous communication of the heat shock response. Considering the aging-related decline of function in the neuronal HSR and its relationship to protein aggregation, manipulation of glial *hsf-1* emerges as a promising tool to tackle aging and neurodegenerative phenotypes broadly.

## Methods

Thermotolerance-Worms were synchronized by bleaching as described here, L1 arrested, and plated on HT115 bacteria. At late D1, 15 worms per plate with 5 plates per condition were exposed to 34ºC heat via incubator for 13-16 hours. Plates were then removed from the incubator and manually assessed for movement and pharyngeal pumping, using light head taps where necessary, to determine survival. Worms that displayed internal hatching or crawled onto the side of the plate and desiccated were censored and omitted from the final analysis. Percent alive was calculated using the number of living worms divided by the total number of worms less censored worms for each strain.

Lifespan-Lifespans were performed as previously described (*34*). In brief, worms were synchronized by bleaching, L1 arrested, and plated on HT115 bacteria. On Day 1 of adulthood, worms were moved to fresh plates with 15 worms per plate and 10 plates per condition. Living worms were counted every day, and occasionally every other day, for the duration of the lifespan. Life was assessed by movement, pharyngeal pumping, or response to a light head touch. Worms were censored if they crawled onto the side of the plate and desiccated, if they displayed internal hatching, or had extruded vulvas/intestines.

### Pseudomonas aeruginosa survival

PA14 bacteria was cultured overnight at 37C protected from light in KB media. 20uL was spread onto slow killing plates, which were incubated 24h at 37C protected from light. After plates returned to room temperature, synchronized L4 worms were added to the plates, using 6 plates of 20 worms per plate. Survival was assessed as described above. Missing worms and those crawling onto the side of the plate were censored and omitted from analysis, but bagged worms were counted as dead for this assay. Worms were counted at least once per day, but more frequently near peak death. Statistics were performed as described for lifespan experiments.

### Imaging

Worms were anesthetized using 100uM sodium azide solution on NGM plates immediately, aligned with a worm pick head to tail, and imaged. Fluorescent and brightfield images were collected via the Echo Revolve Microscope. Exposure time and laser intensity were matched within experiments.

### Worm growth and maintenance

Worms were maintained at 15C on NGM plates spotted with 200uL of OP50 bacteria. Worms were chunked or picked for experiments onto NGM plates with 1mL OP50 and grown up at 20C. They were then synchronized for experiments as described here.

### Synchronization

Worms were synchronized by bleaching as previously described(*44*). In brief, worms were collected off plates into 15mL conical tubes using M9 solution. Bleach solution was added until animals dissolved, and the worms were spun down (30s at 1000 RCF) and washed five or more times with M9 before L1 arrest. L1 arrest was performed by suspending worms in M9 in 15mL conical tubes and rotating overnight at 20C before plating on OP50 or HT115 bacteria.

### RNAi feeding

RNAi feeding was performed as previously described (*3*, *34*).

RNA isolation, library preparation, and sequencing: Animals were bleach synchronized and grown to the L4 stage HT115 plates. At least 2,000 animals per condition per replicate were washed off plates using M9 and collected. After a 30 second spin at 1,000 RCF, M9 was aspirated, replaced with 1mL Trizol, and the tube was immediately frozen in liquid nitrogen to be stored at −80C for downstream processing. RNA was harvested after 3 freeze thaw cycles in liquid nitrogen/37C water bath. After the final thaw, 200uL (1:5 chloroform: Trizol) of chloroform solution were added to the sample, vortexed, and the aqueous phase was collected after centrifugation in a gel phase lock tube. RNA was isolated from the obtained aqueous phase using a Qiagen RNeasy MiniKit according to manufacturer's directions. Library preparation was performed by Azenta Genewiz as follows: Extracted RNA samples were quantified using Qubit 2.0 Fluorometer (Life Technologies, Carlsbad, CA, USA) and RNA integrity was checked using Agilent TapeStation 4200 (Agilent Technologies, Palo Alto, CA, USA). RNA sequencing libraries were prepared using the NEBNext Ultra RNA Library Prep Kit for Illumina following manufacturer’s instructions (NEB, Ipswich, MA, USA). Briefly, mRNAs were first enriched with Oligo(dT) beads. Enriched mRNAs were fragmented for 15 minutes at 94 °C. First strand and second strand cDNAs were subsequently synthesized. cDNA fragments were end repaired and adenylated at 3’ends, and universal adapters were ligated to cDNA fragments, followed by index addition and library enrichment by limited-cycle PCR. The sequencing libraries were validated on the Agilent TapeStation (Agilent Technologies, Palo Alto, CA, USA), and quantified by using Qubit 2.0 Fluorometer (Invitrogen, Carlsbad, CA) as well as by quantitative PCR (KAPA Biosystems, Wilmington, MA, USA). The sequencing libraries were clustered on 1 lane of a flow cell. After clustering, the flow cell was loaded on the Illumina HiSeq instrument (4000 or equivalent) according to manufacturer’s instructions. The samples were sequenced using a 2×150bp Paired End (PE) configuration. Image analysis and base calling were conducted by the HiSeq Control Software (HCS). Raw sequence data (.bcl files) generated from Illumina HiSeq was converted into fastq files and de-multiplexed using Illumina's bcl2fastq 2.17 software. One mismatch was allowed for index sequence identification.

### RNA sequencing analysis

For RNA-seq analysis, the sequencing data were uploaded to the Galaxy project web platform and the public server at *https://usegalaxy.org* was used to analyze the data (*45*). Paired end reads were aligned using the Kallisto quant tool (Version 0.46.0) with WBcel235 as the reference genome. Fold changes and statistics were generated using the DESeq2 tool with Kallisto quant count files as the input. Volcano plots were generated using GraphPad Prism software (Version 9.2.0 (283)) on the fold change and adjusted-p values generated by the previous analysis. GO terms for differentially expressed genes were analyzed by using the GOrilla tool (http://cbl-gorilla.cs.technion.ac.il/#ref) on lists of genes that were up or down regulated (Log2FC > 1, <1, respectively) with an adjusted P-value <= 0.05 (*29*, *46*). The raw RNA-seq data were uploaded to the NCBI short read archive (PRJNA801195). Access for reviewers is available at https://dataview.ncbi.nlm.nih.gov/object/PRJNA801195?reviewer=9jun897nhgr342vonpa0v3a49p.

### Generation and integration of arrays

The *hlh-17* promoter was cloned into a vector containing full length *hsf-1*, with sequences as previously described (*3*, *17*). Wild type N2 strain worms were injected with the *hlh17p::hsf-*1; *unc-54* 3’UTR plasmid and *myo-2p::tdtomato* co-injection marker. Integration of extrachromosomal array lines was performed by gamma irradiation (*Is2(hlh-17p::hsf-1)*) or by UV irradiation (*Is1(hlh-17p::hsf-1)*). Integrated lines were then backcrossed at least eight times to the wild type N2 strain.

### Brood size

Synchronized L4 animals were picked individually onto fresh HT115 bacteria plates and allowed to lay eggs for 24h at 20C. They were then moved to fresh plates for each consecutive 24h period for the duration of the reproductive lifespan for at least five days. Progeny plates were allowed to grow up at 20C for two days, and surviving larvae were imaged using the MBF Bioscience WormLab imaging system and counted.

### Heat shock for imaging

Synchronized worms were placed in a 34C incubator for 2h, followed by a recovery for 2h at 20C, at which time worms were imaged or sorted as described.

Prediction of HSF-1 binding sites in *C. elegans* promoters: HSF-1 binding sites were predicted in the upstream regions of coding sequences using the FIMO tool (version 5.0.5) on MEME Suite (*47*, *48*). Briefly, 500bp upstream flanks of all annotated coding genes were downloaded from the WormBase ParaSite to represent putative promoter regions (*49*). The HSF1 position weight matrix (PWM) was downloaded from JASPAR (matrix ID MA0486.2) (*50*). FiMO was run with the HSF1 PWM as input motif and the putative promoter regions as input sequences with a match P-value < 1E-5 to find 646 genes with HSF-1 binding sites. GO term analysis using the GOrilla tool confirmed the top GO terms of these genes to include chaperone mediated folding (GO:0061077), protein folding (GO:0006457) and response to heat (GO:009408).

### COPAS biosorting and analysis

Worm sorting using the COPAS biosorter (Union Biometrica) was performed as previously described (*44*). In brief, worms were heat shocked as described for *hsp-16.2p::GFP* conditions. Then, they were washed off plates into the sample cup using M9 and sorted. Laser PMT values were consistent within experiments. All raw data was saved. For analysis, reads with time of flight (TOF) greater than 100 and extension (EXT) greater than 50 were included and reads with lower values were excluded. Reads for which EXT or green peak height reached the maximum (saturated) value of 65532 were excluded. Normalized fluorescence was calculated by dividing green peak height by TOF. For red-headed animals, worms with a red peak height of 1000 or greater were included, and lower values were presumed array negative and were excluded.

### Genetic crosses

Males were generated either by heat exposure or by crossing to wild type males. Hermaphrodites and males of interest were placed on NGM plates with a small amount of OP50 bacteria and allowed to mate. Progeny were singled onto individual plates for the F1 and the subsequent F2 generation and were screened for relevant phenotypes.

### Statistical analyses

Statistical analysis was performed using GraphPad Prism 9.2.0(283), excepting RNA sequencing analysis which was performed as described above. Individual analyses are as described in figure legends. Lifespans were analyzed by Gehan-Breslow-Wilcoxon test. Two condition comparisons were otherwise analyzed by two-tailed t test, with Welch’s correction where applicable, and more than two condition comparisons were analyzed by one-way ANOVA with Sidak’s multiple comparisons.

## Author Contributions

HKG and AD conceived of the study, and HKG designed experiments. HKG, PAF, SUT, CLP, HC, TDH, SSC, and ES performed experiments. HKG, PAF, SUT, and AEF generated strains and constructs. HKG and PAF analyzed data and designed figures. SUT also contributed artistic work. HKG wrote the manuscript. All authors reviewed and edited the manuscript.

## Acknowledgements

We thank Dr. Koning Shen and Melissa Metcalf with all Dillin lab members for helpful comments and conversations throughout the project. Some strains were provided by the CGC, which is funded by NIH Office of Research Infrastructure Programs (P40 OD010440). HKG is supported by NSF Grant # DGE175814 and NIA 1F99AG068343-01. CLP is supported by 5F32AG065381. AD and the lab are supported by the Howard Hughes Medical Institute, NIEHS R01ES021667, and NIA R01AG059566.

## Figure legends

Supplementary Figure 1: Phenotype of CEPsh glial *hsf-1* and reproductive effects A) Lifespan of *Is2(hlh-17p::hsf-1)* is increased relative to wild type N2 animals. Median of N2 = 21 days, median of *Is2(hlh-17p::hsf-1)* = 25 days, p < 0.001. B) Total brood size over the reproductive lifespan; *Ex(hlh-17p::hsf-1)* have significantly fewer progeny than wild type N2, p < 0.0001. Error bars are SD C) Brood size depicted across days of the reproductive lifespan. Error bars are SD D) Thermotolerance of *Is2(hlh-17p::hsf-1)* is increased relative to wild type N2 animals, p= 0.03. *hsp-16.2p::GFP* transcriptional reporter worms *Is2(hlh-17p::hsf-1)* versus wild type N2 after mild heat stress and recovery, lined up head to tail. F) *hsp-70p::GFP* transcriptional reporter worms *Is2(hlh-17p::hsf-1)* versus wild type N2 after mild heat stress and recovery, lined up head to tail.

Supplementary Figure 2: Thermotolerance of neurotransmitter mutants. Mutants for each neurotransmitter alone are displayed in blue, and CEPsh glial *hsf-1* with the relevant neurotransmitter mutant is displayed in purple. A) Thermotolerance of *Ex(hlh-17p::hsf-1); cat-2(n4547)* is significantly increased relative to *cat-2(n4547)* alone, p= 0.03. B) Thermotolerance of *Ex(hlh-17p::hsf-1); tdc-1(n3419)* is significantly increased relative to *tdc-1(n3419)* alone, p= 0.04. C) Thermotolerance of *Ex(hlh-17p::hsf-1); unc-25(e156)* is significantly increased relative to *unc-25(e156)* alone, p=0.02. D) Thermotolerance of *Ex(hlh-17p::hsf-1); eat-4(ky5)* is not significantly increased relative to *eat-4(ky5)* alone, though there is a trend towards increase. E) Thermotolerance of *Ex(hlh-17p::hsf-1); unc-17(e245)* is not significantly increased relative to *unc-17(e245)* alone, though there is a trend towards increase.

Supplementary Figure 3: Thermotolerance of CEPsh glial *hsf-1* on RNAis. A) Thermotolerance of *Ex(hlh-17p::hsf-1)* eating *hsf-1* RNAi bacteria (blue) is not significantly increased relative to that of wild type N2 worms eating *hsf-1* RNAi (purple). B) Thermotolerance of *Ex(hlh-17p::hsf-1)* eating *daf-16* RNAi bacteria (blue) is not significantly increased relative to that of wild type N2 worms eating *daf-16* RNAi (purple).

## Notes

https://dataview.ncbi.nlm.nih.gov/object/PRJNA801195?reviewer=9jun897nhgr342vonpa0v3a49p

### Competing Interest Statement

The authors have declared no competing interest.

