## Supplementary figures and images for "Glia of *C. elegans* coordinate the heat shock response independent of the neuronal thermosensory circuit and serotonin"

### Supplemental Figures

Supplementary Figure 1

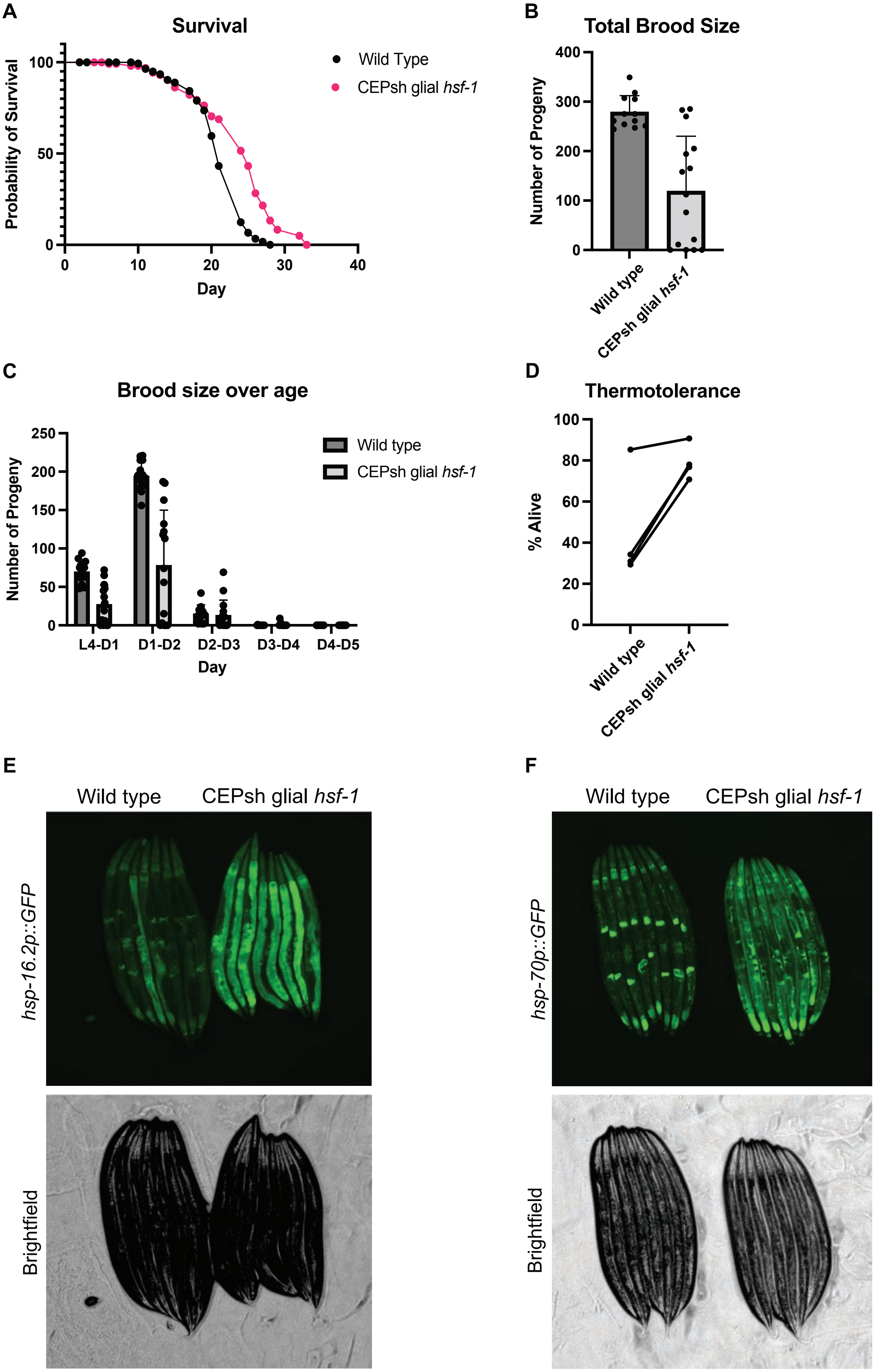

#### Supplementary Figure 2

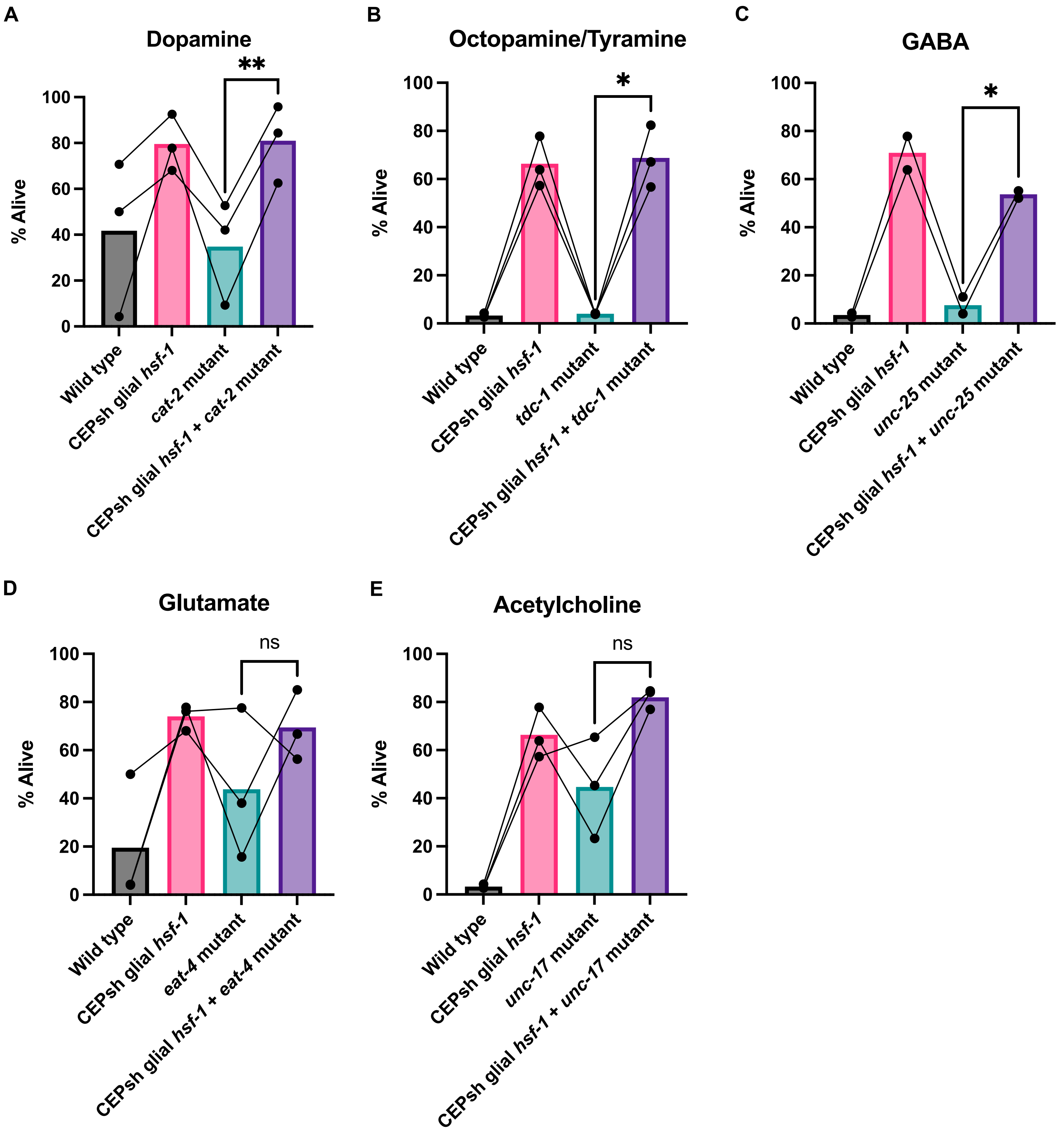

**Supplementary Figure 3**

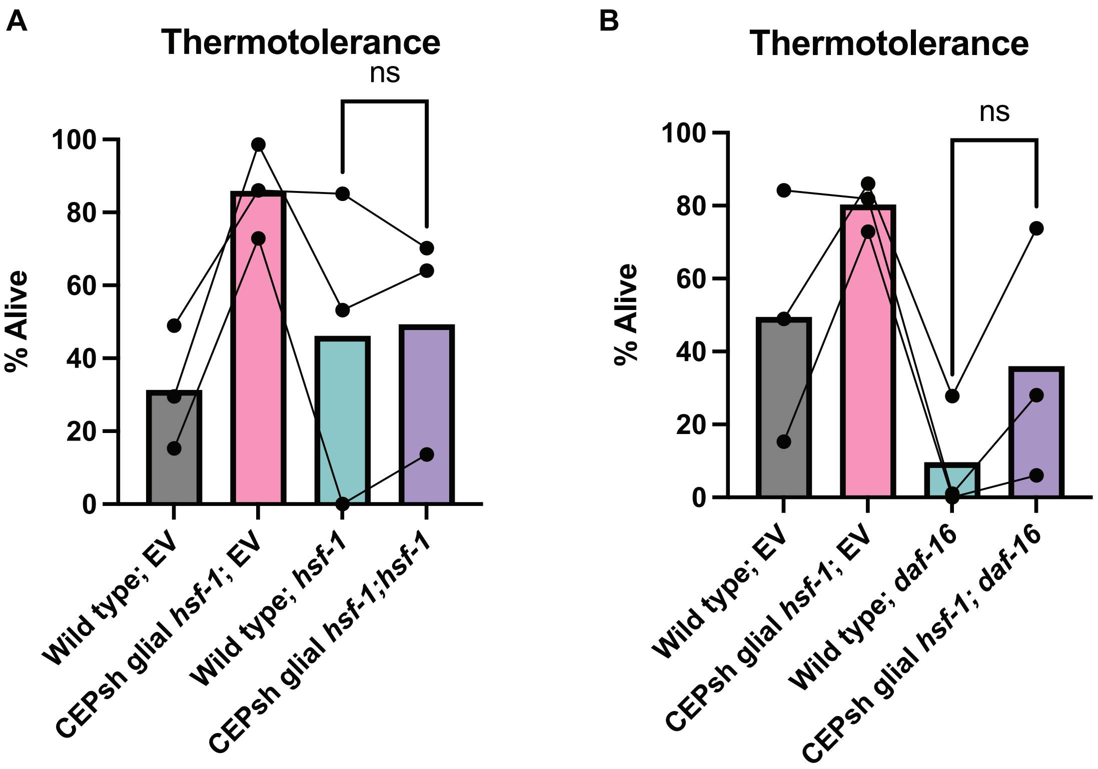
